# Postural sway is not affected by estrogen fluctuations during the menstrual cycle

**DOI:** 10.1101/2022.07.18.500469

**Authors:** Sasha Reschechtko, Thuy Ngoc Nguyen, Michelle Tsang, Kristine Giltvedt, Mark Kern, Shirin Hooshmand

## Abstract

When people stand still, they exhibit a phenomenon called postural sway, or spontaneous movement of the body’s center of pressure, which is related to balance control. In general females show less sway than males, but this difference only begins to appear around puberty, pointing to different levels of sex hormones as one potential mechanism for sway sex differences. In this study, we followed one group of young females using oral contraceptives (n = 32) and one group not using oral contraceptives (n = 19), to investigate associations between estrogen availability and postural sway, both over the course of the menstrual cycle (in the group not using oral contraceptives) and long-term (between the two groups). All participants visited the lab four times over the putative 28-day menstrual cycle. At each visit, we performed blood draws to measure plasma estrogen (estradiol) levels, and tests of postural sway using a force plate. Due to the hormone-stabilizing effects of oral contraceptives, estradiol levels were higher in participants not using oral contraceptives (690.45 pmol/L versus 464.50 pmol/L), reflecting higher estradiol concentrations during the late follicular and luteal phase. Postural sway was similar on average for participants not using oral contraceptives (21.47 cm versus 23.56 cm). Overall, we found no significant effects of the estimated menstrual cycle phase – or absolute levels of estradiol – on postural sway.

## Introduction

Estrogen plays a variety of roles in extragonadal tissues. In addition to its wide-ranging effects on bone (Cutler, 1997; Kalervo Väänänen & Härkönen, 1996; Riggs, 2000), estrogen plays a role in the formation, maintenance, and function of muscle, connective tissue, and neural tissue (Chidi-Ogbolu & Baar, 2019; Cui et al., 2013; Hansen, 2018; Leblanc et al., 2017). During the menstrual cycle estrogen levels can vary by orders of magnitude (Buchanan et al., 1986; Denver et al., 2019). Despite these large fluctuations in estrogen levels and estrogen’s widespread effects, it remains unclear whether estrogen availability affects motor behaviors or performance.

One important motor ability is balance, or our ability to maintain an upright posture without falling over. Postural sway – a measure of spontaneous shifts of a person’s center of mass as they stand stationary – is consistently lower in females compared to males, but this difference does not appear until around puberty and is not accounted for by other likely covariates like height or weight (Goble & Baweja, 2018a, 2018b; Moran et al., 2020). While postural sway is not the same as balance and may reflect multiple neural processes (Carpenter et al., 2010; Latash & Zatsiorsky, 2016; Mochizuki et al., 2006), it is often interpreted as related to balance in that people who show more postural sway are considered to have worse balance and be more likely to fall (Fernie et al., 1982; Johansson et al., 2017).

Balance depends on both sensory and motor factors (Mancini & Horak, 2011; Shumway-Cook & Horak, 1986). Estrogen can alter the structural properties of connective tissues by changing the amount of collagen incorporated into them, the formation of collagen-elastin crossbridge structures, and connective tissue metabolism (Chidi-Ogbolu & Baar, 2019; Hansen, 2018; Leblanc et al., 2017). These changes could affect the mechanical connections between sensory receptors and muscles, but observations of functional changes due to altered mechanical properties in vivo during the menstrual cycle do not show a consensus. While one group has consistently indicated changes in connective tissue and muscle properties over the menstrual cycle (Lee et al., 2013; Petrofsky & Lee, 2015; Yim et al., 2018), others have not reported functional changes on that timescale (Bryant et al., 2008; Burgess et al., 2009; Ericksen & Gribble, 2012; Kubo et al., 2009).

A number of studies have reported that balance-related behavioral measures change over the course of the menstrual cycle (Darlington et al., 2001; Maged et al., 2017; Mokošáková et al., 2018; Petrofsky & Lee, 2015; Sung & Kim, 2018). In conjunction with observations about the timing and prevalence of certain knee and ankle injuries (Wojtys et al., 2002), these findings have been interpreted as indicative of increased injury risk and been taken as evidence supporting the prophylactic use of oral contraceptives to decrease musculoskeletal injury risk in female athletes. However, a number of the cited studies use small participant cohorts (≤ 15: (Darlington et al., 2001; Fridén et al., 2003; Mokošáková et al., 2018; Petrofsky & Lee, 2015) and some use measures of balance that are difficult to replicate or interpret because they are related to proprietary testing apparatus instead of physical measures of sway (Darlington et al., 2001; Fridén et al., 2003; Maged et al., 2017; Yim et al., 2018). Finally, these studies rarely measure serum hormone levels to investigate hormone level as a mechanism. Here we measure both postural sway and blood hormone levels four times over the course of the menstrual cycle in 19 young female participants who reported natural menstrual cycling as well as 32 young females taking oral contraceptives.

## Methods

### Participants

We recruited healthy young female participants who were either using oral contraceptives or not using oral contraceptives to participate in this study. Participants were required to be between 18-25 years old, have a BMI between 18-32 kg/m^2^, be non-smoking, consume fewer than 2 alcoholic drinks per day and not be pregnant or lactating. Participants using oral contraceptives were required to have been using for at least one year and no more than 5 years; individuals using other forms of hormonal contraceptives were not eligible to participate. Potential participants were given a questionnaire regarding their menstrual history and oral contraceptive use; based on their answers, participants were divided into a group using oral contraceptives and a group who were not using oral contraceptives (naturally cycling). These participants were part of a larger study investigating the effects of oral contraceptives on bone health. Naturally cycling participants had not been using hormonal contraceptives for at least 3 months before they participated. We conducted this study in accordance with procedures approved by the San Diego State Institutional Review Board’s Human Research Protection Program.

Following data collection, we imposed additional inclusion criteria on participants who were not using oral contraceptives to ensure they showed a typical hormonal profile during the menstrual cycle. We only included participants who (1) showed peak progesterone levels at Visit 4 (mid luteal phase), and (2) showed an increase in estradiol levels from Visit 1 (early follicular phase) to Visit 3 (late follicular phase). We imposed these criteria to ensure a participants showed relatively regular menstrual cycle; it has been suggested that a major source of variability in previous literature investigating the menstrual cycle is the inclusion of females with atypical menstrual cycles in supposedly eumenorrheic groups (Dam et al., 2022).

### Testing protocol

Following an initial lab visit for familiarization and screening, we scheduled each participant to make four visits for blood collection and balance testing. For participants who were not using oral contraceptives, these visits corresponded to menstrual cycle days 2, 4, 11, and 21 (first day of follicular phase, early follicular phase, ovulation, and mid luteal phase, respectively). For participants using oral contraceptives, these visits corresponded to pill pack days 21 (start of placebo pills), 24, 3, and 13. We chose these intervals for participants on oral contraceptives to coincide with the start of the estrogen-free pill phase of the oral contraceptives (day 21) and to match the time between visits in the naturally cycling group. Participants always visited the lab in the same order: their first testing visit corresponded to cycle day 2 (21 for oral contraceptive users) – participants contacted the study group to schedule their first test as soon as menses began – and the last visit was day 21 (13 for oral contraceptive users). The balance testing paradigm we used is unaffected by practice or prior exposure effects (Hearn et al., 2018), so there should not be systematic changes due to the visit sequence itself. We asked participants to fast for 10 hours before their scheduled visits and we instructed oral contraceptive users who consumed their pill in the morning to wait until after the blood draws. We centrifuged blood samples immediately after blood draws for 15 minutes at 4000 x g at 4° C and subsequently stored the samples at −80° C until analysis.

### Evaluating balance

We evaluated participants’ balance using the BtrackS Balance Test (BBT; Balance Track Systems, Inc. San Diego, CA, USA). The BtrackS system incorporates a standardized force plate and data collection protocol to quantify postural sway during quiet standing. Postural sway results from the natural, spontaneous motion of a person’s center of mass as they stand stationary and reflects the combination of neural outputs and sensory inputs that govern balance. Balance is interpreted as inversely related to postural sway: a person who displays more postural sway (meaning their center of mass moves more) is considered to have worse balance.

The BBT testing protocol involves recording postural sway during three, 20 s trials following one familiarization trial. During these trials, participants are instructed to stand as still as possible on the BtrackS force plate with their eyes closed and feet shoulder width apart. The BtrackS force plate records the center of pressure (the projection of the center of mass onto the ground) at a rate of 25 Hz. Results from the BBT have been validated against laboratory-quality force plates (Goble et al., 2018; O’Connor et al., 2016; Richmond et al., 2018). The outcome measure of the BBT (“BBT Score”) is the average length, in cm, of the path that the center of pressure covers during the 3 trials. While the precise link between balance and postural sway is unknown (Latash & Zatsiorsky, 2016), extensive normative data are available for BBT scores.

### Measuring sex hormones

We measured estradiol, which is the most biologically active and most prevalent form of estrogen, as well as progesterone. We analyzed the level of estradiol in plasma in each sample using ultra-sensitive estradiol ELISA kits (ALPCO, Salem, NH, USA). These ultra-sensitive ELISA kits have a sensitivity of 5.14 pmol/L with a range up to 734 pmol/L. For samples that exceeded the upper limit of the ultra-sensitive ELISA kits, we then used standard ELISA kits (ALPCO, Salem, NH, USA) which have a sensitivity of 36.71 pmol/L with a range up to 11747 pmol/L. To further analyze participants’ hormone profiles, we also measured serum progesterone levels. The ELISA kits (ALPCO, Salem, NH, USA) we used to measure plasma progesterone concentrations have a sensitivity of 0.32 nmol/L and a range up to 191 nmol/L.

### Statistical analysis

To test the effect of the menstrual cycle and oral contraceptive use on estradiol levels, we used linear mixed-effects models. We log-transformed estradiol level for normality, as assessed visually via QQ-plots. We used the same methods to assess the effect of the menstrual cycle and oral contraceptive use on the BBT score to investigate postural sway. We report Bayes Factors (BF10) for additional interpretive value; unlike large P-values, small Bayes Factors can be interpreted as evidence for the null hypothesis (Kass & Raftery, 1995), and large Bayes Factors can also indicate that a study is properly powered. These tests use day of menstrual cycle as a proxy for estradiol level, so we also directly investigated whether there was an association between estradiol level and balance by running repeated measures linear correlation on estradiol level and BBT score.

We used JASP (version 0.16.3, JASP Team, 2022) to perform our statistical analyses. We analyzed the effect of the day of the menstrual cycle on hormone levels and postural sway using linear mixed-model analysis. We constructed the linear mixed-effects models using the Linear Mixed Models workflow in JASP with Visit number (4 levels) and Contraceptive status (2 levels) treated as fixed effects and individual participants as random factors for which we estimated random intercepts but not slopes. We performed the same analyses in a Bayesian framework with the Bayesian ANOVA workflow in JASP treating individual participants as random factors and default assumptions. We performed repeated measures correlations using the *rm_corr* command in the pingouin package (Vallat, 2018) for Python.

## Results

### Participant retention and exclusion

In total, 77 female participants completed four sets of balance tests; however, only 58 of the participants completed full sets of both balance tests and blood draws (32 using oral contraceptives and 26 not using oral contraceptives). This occurred because some participants had not yet fully healed from their first blood draw when they returned for their second blood draw (two days later) and were therefore unable to undergo that blood draw. After imposing additional constraints on hormonal profile in participants who were not using oral contraceptives (described later), we included 19 of 26 participants not using oral contraceptives in our analyses. Physical characteristics and demographic information of the participants we included in analysis are provided in Table 1, and a visualization of testing schedule and participant inclusion is shown in Figure 1.

**Table 1.**

|  |  | *No Contraceptive (n = 19)* | *Hormonal Contraceptive (n = 32)* |
| --- | --- | --- | --- |
| *Age (yr)* | *Mean* | 21.89 | 21.34 |
|  | *SD* | 2.08 | 1.73 |
| *Height (cm)* | *Mean* | 162.18 | 163.78 |
|  | *SD* | 6.02 | 5.82 |
| *Weight (kg)* | *Mean* | 62.27 | 61.02 |
|  | *SD* | 5.82 | 7.65 |
| *Demographics* | *White* | 10 | 20 |
|  | *Hispanic* | 7 | 4 |
|  | *Asian* | 2 | 4 |
|  | *Black* | 0 | 2 |
|  | *Other* | 0 | 2 |

**Figure 1.**
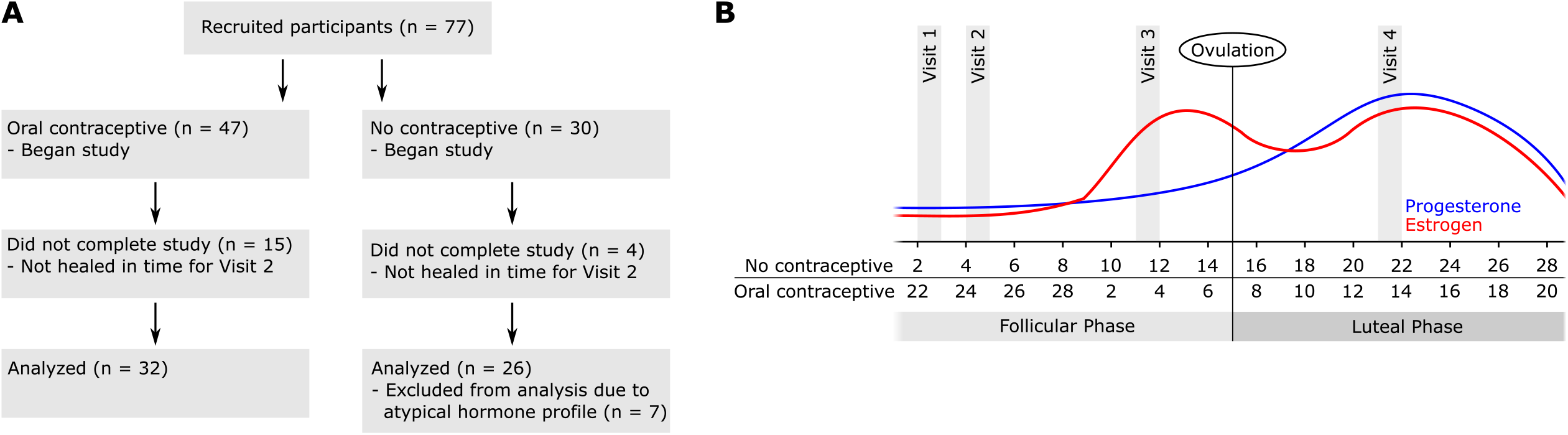
Experimental Design and Participant Inclusion. Panel A: workflow indicating participants considered for analysis and exclusion criteria. From 77 participants with full balance data, we included 51 participants in the analyses here. Panel B: schedule of visits and “typical” menstrual cycle hormonal profile adapted from (Reed & Carr, 2000). Testing intervals were the same for participants using oral contraceptives and those not using oral contraceptives, but for those using oral contraceptives testing was chosen to start on the first day pill pack placebo.

### Confirmation of altered estradiol levels due to oral contraceptive use

We observed a large range of estradiol levels in both groups: across all visits and participants, estradiol levels in participants using oral contraceptives ranged from 0.50 pmol/L to 4837.35 pmol/L; in participants not using oral contraceptives, estradiol levels ranged from 19.64 pmol/L to 5685.12 pmol/L. On average across all visits, estradiol levels were higher in the group not using oral contraceptives (690.47 pmol/L) than in the group using oral contraceptives (464.50 pmol/L). These levels are similar to those reported in previous studies (Bryant et al., 2008; Buchanan et al., 1986; Denver et al., 2019). We found a significant interaction between day of menstrual cycle and oral contraceptive use (F_3,174_ = 14.68; P = 2.05e^-8^; BF_10_ = 58.92), indicating that estradiol levels were different between groups during some days of the menstrual cycle. For participants not using oral contraceptives, estradiol levels were higher during days 11 and 21 than days 2 and 4. In contrast, estradiol levels did not vary significantly over the course of the menstrual cycle for participants using oral contraceptives, so estradiol levels in the group using oral contraceptives were lower than in the group not using oral contraceptives during days 11 and 21. These results are shown in Figure 2.

**Figure 2.**
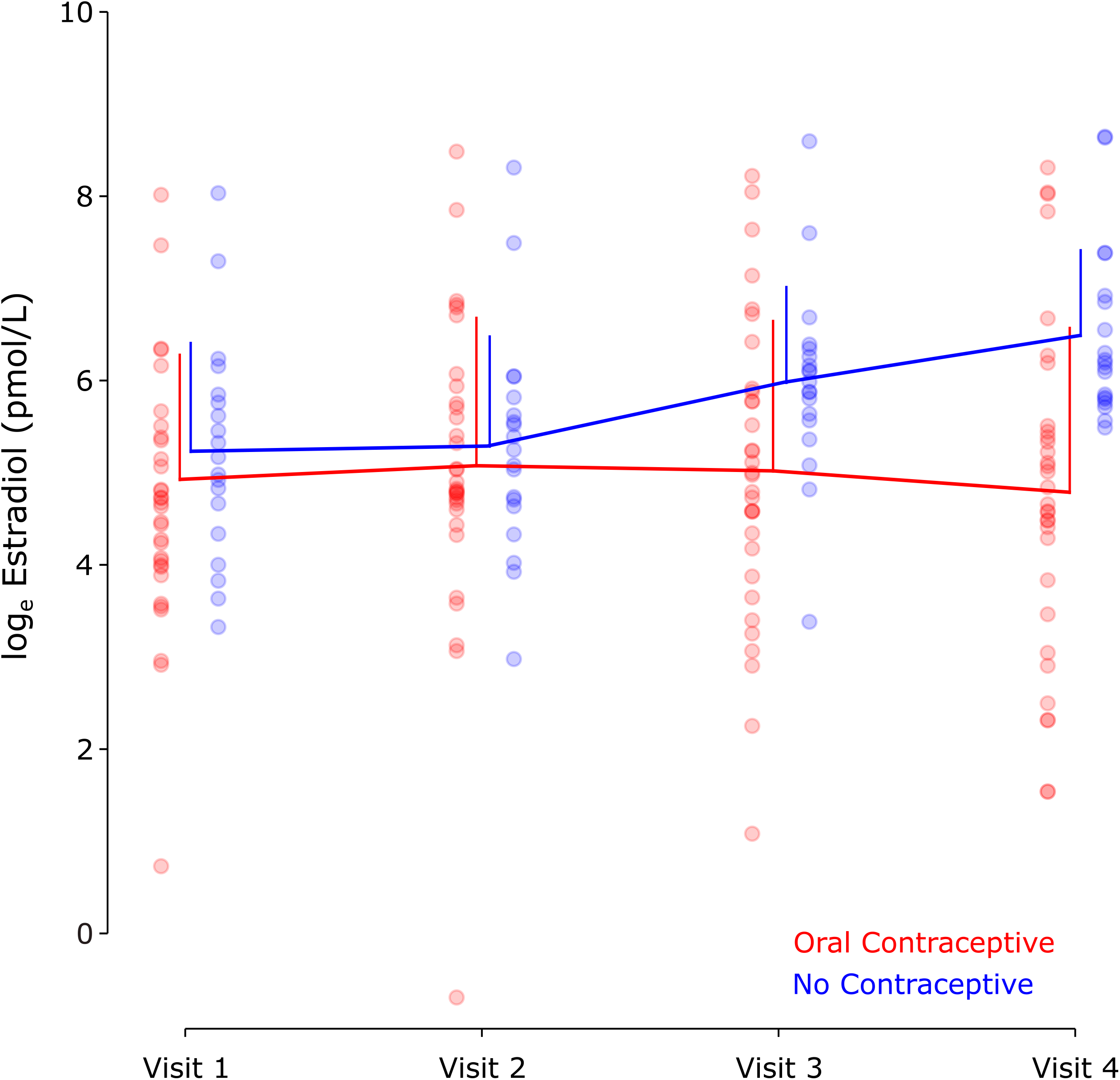
Estradiol concentrations. Plasma estradiol concentrations as measured via blood draw during the four experimental visits. Blue: females not using oral hormonal contraceptives (N = 19); Red: females using oral hormonal contraceptives (N = 32). Each dot represents a single participant. Error bars are SD. Estradiol data are log-transformed for analysis and visualization. In the participant group using oral contraceptives, visit days correspond to days 2, 4, 11, and 21 of the putative 28-day menstrual cycle.

### Postural sway is unaffected by menstrual cycle

Across days and participants, we observed a range of pathlengths from 13.56 cm to 43.64 cm in the participants using oral contraceptives and from 13.66 cm to 32.15 cm in participants not using oral contraceptives. Postural sway for participants using oral contraceptives (23.79, 23.65, 23.2, and 23.6 cm for visits 1-4, respectively) was not significantly different from postural sway in participants not using oral contraceptives (21.31, 20.84, 22.31, and 21.42 cm for visits 1-4) (F_1,49_ = 2.35; P = 0.132; BF_10_ = 0.85), nor did these values vary across day of the menstrual cycle (F_3,147_ = 0.163; P = 0.92; BF_10_ = 0.026). Compared to previously published normative data using this postural sway paradigm, all of these values are between the 30th and 50th percentiles for females ages 15-29 (Goble & Baweja, 2018b). Figure 3 shows the average values and all individual participants’ data, which gives the visual impression that the variability of the data is high compared to mean differences, and the magnitude of the differences is lower than the minimum meaningful difference for this test (5cm; Goble et al. 2016).

**Figure 3.**
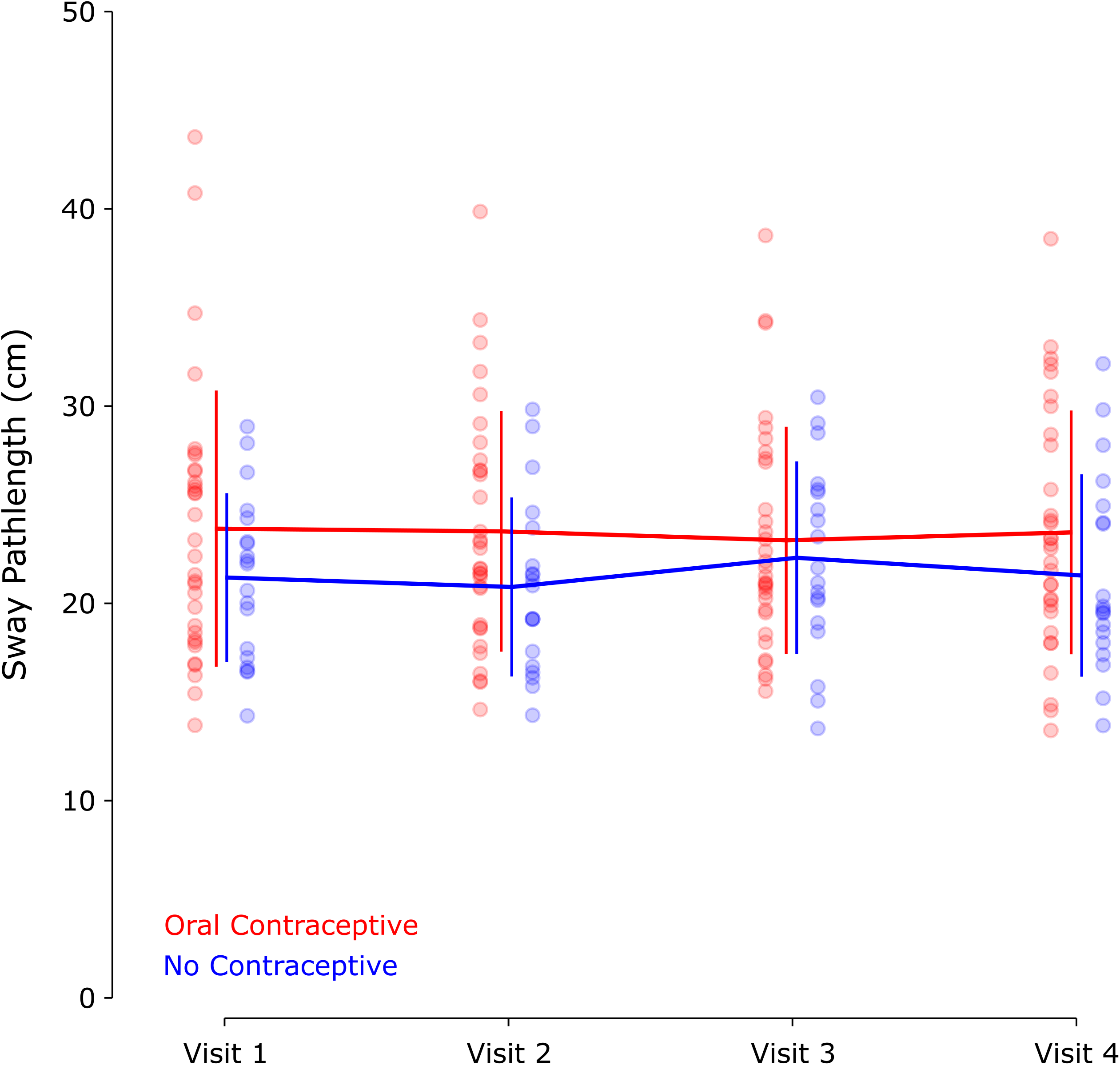
Postural sway. Postural sway, quantified as the average pathlength of the center of pressure during three 20 s trials of eyes-closed quiet standing. Blue: females not using oral contraceptives (N = 19); Red: females using oral contraceptives (N = 32). Error bars are SD.

### Estradiol level is minimally associated with postural sway

Finally, we directly tested whether estrogen level was associated with postural sway using repeated measures correlation (Bakdash & Marusich, 2017; Vallat, 2018) of estradiol level with postural sway. The repeated measures correlation between estradiol concentration and postural sway was not significant for participants using oral contraceptives (r = 0.08; P = 0.46; 95% CI = [-0.12, 0.27]), or for participants not using oral contraceptives (r = 0.10; P = 0.44; 95% CI = [-0.16, 0.35]). These data are shown in Figure 4. Additionally, we compared the two groups in a mixed linear effects model incorporating the actual estradiol concentrations rather than visit days (visit days were used in the previously described analyses). Mean postural sway for participants not using oral contraceptives was 21.27 cm (95% CI = [19.11, 23.44]) and 23.7 cm (95% CI = [22.07, 25.34]) for participants using oral contraceptives. Model slope (indicating the estimated effect of estradiol concentration on postural sway) in participants not using oral contraceptives was 0.40 (95% CI = [-.62, 1.41]) and 0.49 (95% CI = [-.27, 1.25]) in participants using oral contraceptives.

**Figure 4.**
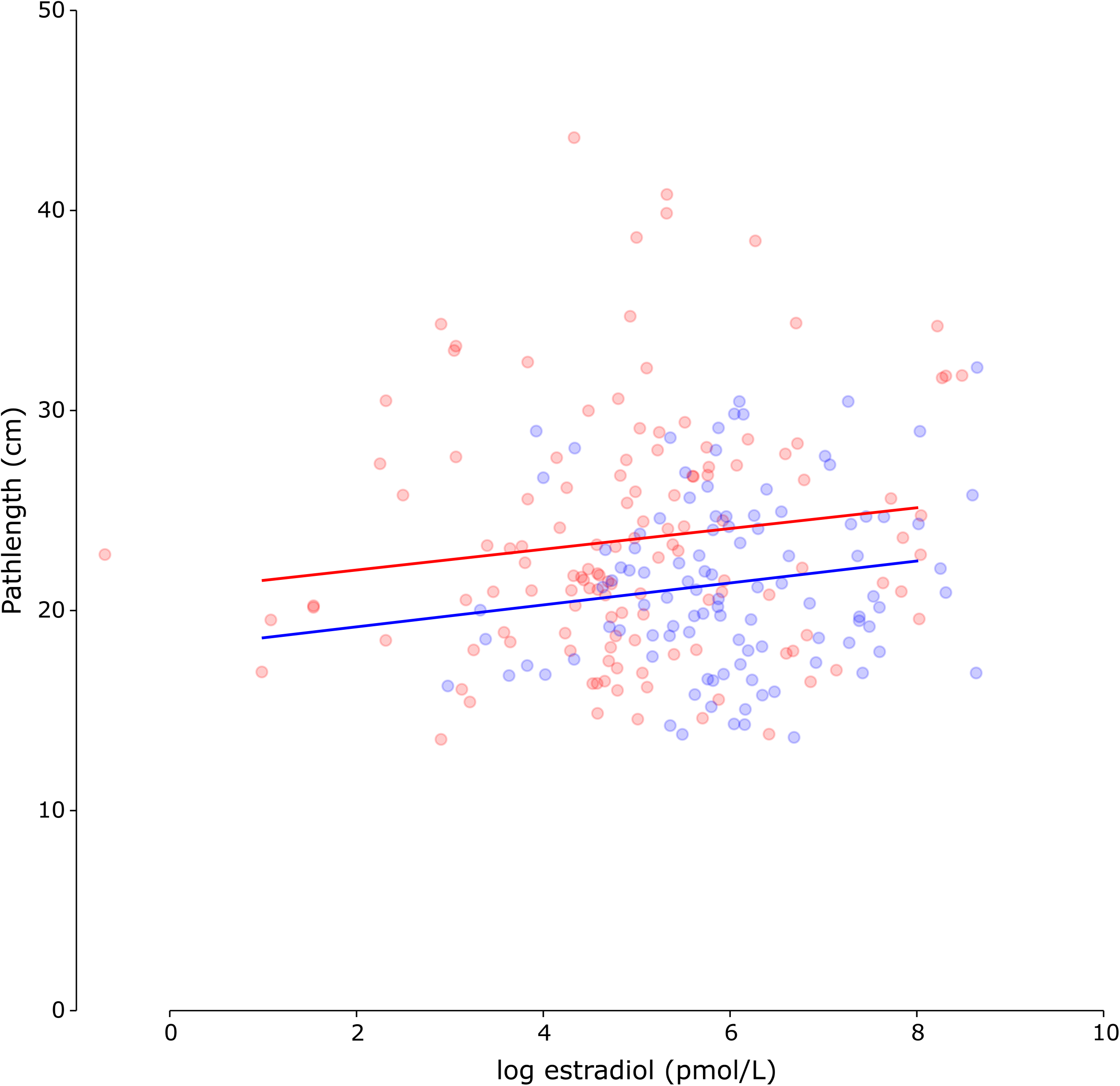
Comparison of estradiol and postural sway. Scatterplot of estradiol concentration (log transformed) and postural sway (pathlength of center of pressure excursion) for females using oral contraceptives (Blue; N = 19) and females not using oral contraceptives (Red; N = 32). Repeated-measures correlations were not significant in either group. Solid lines represent best linear fits.

## Discussion

As expected, our study participants not using oral contraceptives showed changes in estradiol levels during the menstrual cycle, whereas those using oral contraceptives did not. However, our measures of balance do not provide evidence that postural sway changes over the course of the menstrual cycle. Postural sway in participants using oral contraceptives (who had lower levels of estradiol) was not significantly different from that in participants who were not using oral contraceptives, although larger sample size might be required to elucidate this point further (BF_10_ ≈ 1). Further, when we compared estradiol levels to participant sway across all participants, we found no evidence that varying levels of estradiol affect postural sway in people using oral contraceptives.

### Previous association of balance and hormone levels

Previous studies report conflicting findings about whether balance-related measures change over the course of the menstrual cycle. Some groups have reported changes in balance over the course of the menstrual cycle during “more challenging” tests, for example eyes closed in combination with unstable surfaces and tandem foot position in which one foot is in front of the other (Petrofsky & Lee, 2015). While it is possible we would have seen differences if we had used other balance-related testing protocols, there are also studies that report changes in balance metrics using methods that are similar to those we used (eyes closed and standing on a firm surface) over the course of the menstrual cycle (Mokošáková et al., 2018), as well as studies that do not report changes using dynamic balance tasks (Ericksen & Gribble,2012).

When studies find a difference in balance metrics over the course of the menstrual cycle, worse balance is generally reported when estrogen levels are high. Curiously, the direction of these results is counter to the observed sex differences in balance. While females generally show lower postural sway than men, the significant associations reported in literature have indicated that increases in estrogen are associated with increased postural sway; this seems counterintuitive given that men tend to have lower estrogen levels and higher sway. This could have occurred due to misidentification of phases of the menstrual cycle or atypical menstrual cycle in the participants analyzed (Dam et al., 2022) or potentially due to publication bias.

We might have observed significant differences over the course of the menstrual cycle if we had used different balance measures, but we elected to use a well-documented balance test with a large amount of normative data instead. Most previous studies investigating menstrual-cycle related fluctuations in balance control with objective measurement of balance have used custom instrumentation and reported measures in arbitrary units (Darlington et al., 2001; Fridén et al., 2003) or units that are specific to the balance testing apparatus (Maged et al., 2017; Sung & Kim, 2018). This makes it difficult to link experimental findings to any physical quantity to understand behavior. As such, we think that the easily replicable testing paradigm and interpretability of postural sway as a straightforward measure of postural control in the present study are important contributions.

### Different timescales of fluctuations in estrogen levels and motor adaptation

While hormone levels vary cyclically over the course of the menstrual cycle, they also vary over lifetime via aging and with long-term exposure to exogenous hormones, for example via oral contraceptives. This study tested both those short-term fluctuations in the group of participants not using oral contraceptives and longer-term fluctuations between the two participant groups.

Although we did not find changes in postural control in either case, there is stronger evidence against changes due to short term fluctuations according to the day of the menstrual cycle than during long term fluctuations between females on and off oral contraceptives.

Several studies have not found changes in the structural properties of connective tissue over the course of the menstrual cycle (Burgess et al., 2009, 2010; Kubo et al., 2009). However, comparisons of males and females show differences in connective tissue properties (Beynnon et al., 2005; Shafiei et al., 2016), and some similar results are reported when comparing apparently eumenorrheic females and females taking oral contraceptives (Bryant et al., 2008). These latter findings agree with changes that would be expected due to changes in connective tissue metabolism (Chidi-Ogbolu & Baar, 2019; Hansen, 2018; Leblanc et al., 2017) although much of the work in this area has not been performed in humans.

We did not find differences in postural sway related to changes in hormone levels due to oral contraceptive use or phases of the menstrual cycle. Even though the physical properties of tissues change, other types of voluntary behavior like maximum voluntary force production does not change over the course of the menstrual cycle either (Bryant et al., 2008; Elliott et al., 2005). Changes in tissue properties, which may manifest themselves in differences in the effects of muscle activation or sensory inputs, are probably among the factors that the nervous system accommodates over the course of development and aging to support our ability to successfully interact with the world over lifespan. While it may seem surprising that we could adapt to changes in the body on a short timescale like the menstrual cycle, motor adaptation paradigms routinely report adaptation on the timescale of hours (Maeda et al., 2020; Weiler et al., 2019) even in the context of supposedly “hard wired” responses like short-latency reflexes. Our results suggest that postural control could similarly quickly adapt to such systemic changes over the course of days during the menstrual cycle.

### Limitations

Menstrual cycle atypicality is common among females who are not using hormonal contraceptives. Our original study participant pool included participants who may have had luteal phase-deficient or anovulatory cycles, and we subsequently excluded 7 participants who were not using oral contraceptives. While exclusion of these participants did not qualitatively change any statistical outcomes, other factors including the assumption of a putative 28-day menstrual cycle for scheduling participant visits could still have affected our measures, especially as they relate to changes in sway during the menstrual cycle (as opposed to those linked to estradiol concentration). Additionally, we did not record participants’ levels of physical activity, which could also affect menstrual or balance outcomes.

Postural control is difficult to define, and postural sway is not a direct measure of balance. It is possible that our measure of sway did not capture some aspect of balance that is associated with changes in hormone levels. Additionally, in the context of sports injuries, the series of events that lead up to an injury (like a rapid change in direction) may not be closely linked with the factors measured via postural sway. Because of this, it is possible that there are balance- or injury-related measures that would be sensitive to changes in hormone level but which we did not capture using postural sway.

### Concluding Comments

While previous studies have investigated the effects of the menstrual cycle on balance, our study makes the following important contributions. First, we quantified balance using an easily understood measure (postural sway) which is reported in common physical units and can be compared to normative data from more than 16,000 people (Goble & Baweja, 2018b). Second, we recruited a large group of female participants whom we followed to obtain longitudinal data at four specific timepoints related to the menstrual cycle. Finally, and critically, we measured both plasma estradiol concentrations and postural sway in the same participants during the same visits, allowing us to directly investigate the potential relationship between hormone levels and balance, rather than using reported menstrual cycle phase as a proxy for hormone levels which vary widely among individuals.

## Acknowledgements

We gratefully acknowledge the valuable assistance we received from the following students at San Diego State University: Taylor Demasi, Jenna Laughlin, Jeff Moore, and Svitlana Storm.

